# Whole-genome analysis of *de novo* and polymorphic retrotransposon insertions in Autism Spectrum Disorder

**DOI:** 10.1101/2021.01.29.428895

**Authors:** Rebeca Borges-Monroy, Chong Chu, Caroline Dias, Jaejoon Choi, Soohyun Lee, Yue Gao, Taehwan Shin, Peter J. Park, Christopher A. Walsh, Eunjung Alice Lee

**Affiliations:** Division of Genetics and Genomics, Manton Center for Orphan Disease, Boston Children’s Hospital, Boston, MA, USA; Broad Institute of MIT and Harvard, Cambridge, MA 02142, USA; Department of Biomedical Informatics, Harvard Medical School, Boston, MA, USA; Division of Developmental Medicine, Boston Children’s Hospital, Harvard Medical School, Boston, MA, USA; Department of Pediatrics, Harvard Medical School, Boston, MA, USA; Department of Genetics, Harvard Medical School, Boston, MA, USA; Department of Neurology, Harvard Medical School, Boston, MA, USA; Howard Hughes Medical Institute, Boston Children’s Hospital, Boston, MA, USA

## Abstract

Retrotransposons are dynamic forces in evolutionary genomics and have been implicated as causes of Mendelian disease and hereditary cancer, but their role in Autism Spectrum Disorder (ASD) has never been systematically defined. Here, we report 86,154 polymorphic retrotransposon insertions including >60% not previously reported and 158 *de novo* retrotransposition events identified in whole genome sequencing (WGS) data of 2,288 families with ASD from the Simons Simplex Collection (SSC). As expected, the overall burden of *de novo* events was similar between ASD individuals and unaffected siblings, with 1 *de novo* insertion per 29, 104, and 192 births for Alu, L1, and SVA respectively, and 1 *de novo* insertion per 20 births total, while the location of transposon insertions differed between ASD and unaffected individuals. ASD cases showed more *de novo* L1 insertions than expected in ASD genes, and we also found *de novo* intronic retrotransposition events in known syndromic ASD genes in affected individuals but not in controls. Additionally, we observed exonic insertions in genes with a high probability of being loss-of-function intolerant, including a likely causative exonic insertion in *CSDE1*, only in ASD individuals. Although *de novo* retrotransposition occurs less frequently than single nucleotide and copy number variants, these findings suggest a modest, but important, impact of intronic and exonic retrotransposition mutations in ASD and highlight the utility of developing specific bioinformatic tools for high-throughput detection of transposable element insertions.

## Main Text

Retrotransposons contribute to genomic and transcriptomic variability in humans and cause a variety of human diseases^1^. Retrotransposons are a class of mobile DNA elements which can copy themselves into RNA and insert themselves into new regions of the genome. This retrotransposition event is estimated to occur in one out of 20-40, 63-270, and 63-916 live births for Alu, LINE-1 (L1), and SVA elements respectively^2–6^. Although the rates are lower than *de novo* rates of single nucleotide variants (SNVs) (44-82 SNVs per birth^7^), transposable element insertions (TEIs) in both exons and non-coding regions can cause diseases by various mechanisms, including disrupting coding sequences, causing deletions, and altering RNA splicing, which can cause frameshifts and loss of function (LoF)^1; 8^. To date, there are more than 100 cases of TEIs causing diseases^1^, including *de novo* insertions in developmental disorders^9^. TEIs in somatic human tissues have also been implicated in complex diseases, such as cancer^10–15^. A landmark study identified a deep intronic SVA insertion causing exon-trapping in a child with Batten disease, resulting in the development of a personalized antisense-oligonucleotide drug to fix the splicing defect^16^. Thus, the identification of TEs is important for increasing genetic diagnoses^17^ but also creates the promise of developing novel therapeutics for specific mutant alleles.

The detection of TEIs in genome sequencing data requires specific pipelines, given their repetitive nature and short read length^10; 18; 19^. These mutations have previously been excluded from most routine genetic diagnoses and studies, including for ASD. Furthermore, accurate estimation of *de novo* TEIs in healthy individuals is important to understand the contribution of *de novo* TEIs in disease cohorts. Initial methods to determine *de novo* rates of TEIs relied on indirect methods which compared two reference genomes, making assumptions regarding the time to the most recent common ancestor between human reference genomes^4^ and human-chimpanzee divergence time^3^. In order to directly determine *de novo* retrotransposition rates, large cohorts are necessary given the infrequency of these events. More recent studies using short-read sequencing technologies have included fewer than 1,000 families each, leading to uncertainties in estimates, especially for SVA insertions^6; 20; 21^. They have also not accounted for the lower sensitivity of detection on TEIs using short read sequencing^19; 22^.

Autism Spectrum Disorder (ASD) is a heterogeneous developmental disorder characterized by communication deficits, impaired social interactions, and repetitive behaviors^23^. 1 in 54 children are diagnosed with ASD in the United States^24^. Although about 50% of the overall heritability of ASD reflects common variation at a population level^25; 26^, rare inherited and *de novo* copy-number variations and single nucleotide variations confer high risk to developing ASD, and drive ASD risk when present in individual children^27–35^. These rare variants are enriched in simplex families, where both parents are unaffected, with *de novo* copy-number variations and single nucleotide variations contributing to 30% of cases in the Simons Simplex Cohort (SSC)^31^. Although recent ASD studies have included TEIs^20–22^, the smaller sample size and the low rates of *de novo* TEIs limited their analyses leaving the role of *de novo* TEIs in both exons and introns in ASD largely unknown.

In this study, we sought to define the role of transposable elements in ASD by analyzing 2,288 families for *de novo* TEIs at whole genome resolution (Figure 1A). We used a computational tool, *xTea* (https://github.com/parklab/xTea), to detect TEIs in high coverage (~40x) whole genome sequencing (WGS) data from the SSC. The analyzed samples consist of ASD families with one affected individual, two unaffected parents, and for 1,860 of these families, one unaffected sibling which was analyzed as the unaffected control. In order to process this massive amount of >9,000 individual whole genomes, we optimized *xTea* for scalability, especially by reducing memory usage and run-time. A dockerized version of xTea was set up and run through an automated workflow management framework, *Tibanna*^36^ on Amazon Web Services. We used an additional *xTea* filter module to reduce false positives in both polymorphic and *de novo* insertions. This module includes TE type specific filters, including a filter for TEIs which fall in reference TEs with a low divergence rate (<=10%) (see Methods for details). *xTea* candidates were classified as “high” or “low” confidence insertions depending on whether enough insertion supporting features were distributed on both sides of the breakpoint. Here, we only included insertions classified as “high confidence”. We further excluded *de novo* candidates which overlapped with reference and known non-reference (KNR) TEIs^5; 37–45^ (see Methods for details).

**Figure 1.**
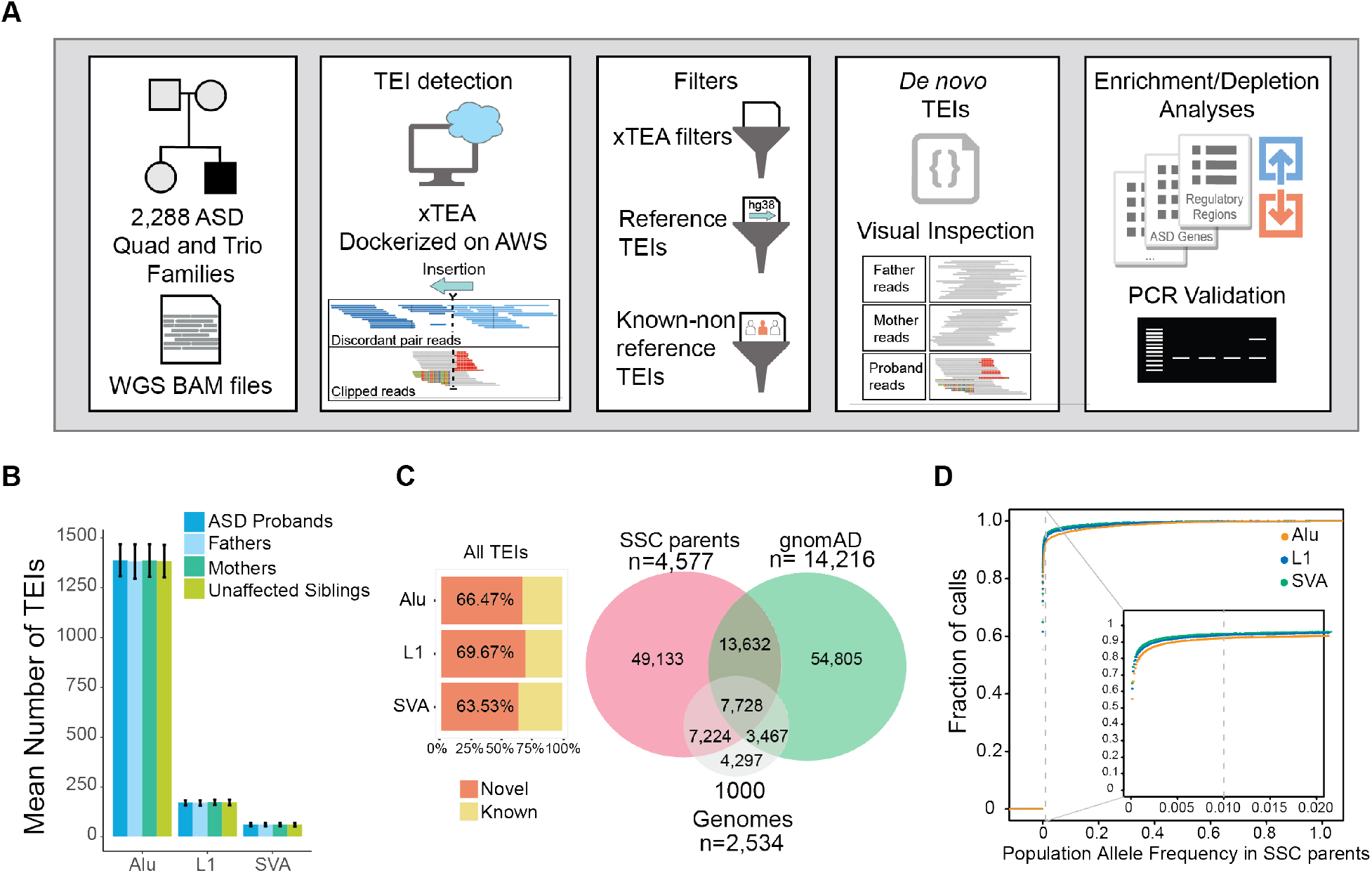
Detection of transposable element insertions (TEIs) in the SSC cohort. **A** Pipeline and analysis overview. Quad and trio bam files were analyzed for TEIs using a dockerized version of *xTea* on the cloud in Amazon Web Services (AWS). Candidate TE insertions were filtered using xTea filters, and filters for regions of the genome with reference and known non-reference TEIs for a high confidence set. A custom pipeline for detection of *de novo* insertions was used, and candidates were manually inspected on the Integrative Genomics Viewer. Enrichment or depletion of TEIs in ASD genes, high pLI genes, genomic regions, and regulatory regions in fetal brain development was tested by simulation analyses. A subset of candidates was validated by full-length PCR. **B** Mean number of TEIs detected in the SSC cohort with standard deviation. **C** Percentage of insertions in the SSC cohort which were not found in previous studies (novel) or overlap with TEIs from previous analyses (known) for all TEIs including those in parents and children (left) and Venn diagram showing overlap with other large cohort studies for TEIs detected in unrelated parental samples in our cohort (right). **D** Cumulative fraction of TEIs in unrelated parental samples which are found at a certain population allele frequency (PAF) within the SSC cohort. 94% L1, 92% Alu, and 95% SVA insertions show <1% PAF.

We detected a total of 86,154 unique polymorphic TEIs (68,643 Alu, 12,076 L1, and 5,435 SVA) in the entire cohort (parents and children) (Figure S1A and Table S1). Each individual genome carried 1,618 polymorphic TEIs on average (1,385 Alu, 172 L1, and 61 SVA) comparable with previous analyses^5; 10; 46^, and the numbers were consistent across different family members (Figure 1B and Figure S1A). 74% of these TEIs (50,507 Alu, 9,247 L1, 4,273 SVA) were observed in either more than one individual in this cohort (71%; 48,189 Alu, 8,821 L1, 4,021 SVA) or in previous studies (33%; 23,018 Alu, 3,663 L1, 1,982 SVA) (Figure S1B), suggesting that the majority of these calls are bona fide. However, more than 60% of calls were novel and had not been detected before in gnomAD^47^ or in the 1000 genomes cohort^44^ (Figure 1C and Figure S2). In 4,577 unrelated parental samples in our cohort we detected 77,717 TEIs, compared to the 79,632 insertions detected from 54,805 individuals in the gnomAD-SV cohort^47^. Additionally, insertions in our cohort had a higher overlap with previously published insertions from 2,534 individuals in the 1000 genomes cohort^44^ (Figure 1C). The majority of parental TEIs were rare, for example, >92% of TEIs having <1% population allele frequency (PAF) within the analyzed cohort (Figure 1D and Figure S3), which is similar to previous findings of structural variants^47^.

We identified 158 *de novo* TEIs from all children (Table S2), which did not have supporting reads in parental raw *xTea* output files and were confirmed via manual inspection on IGV^48^. Previous studies have generally reported *de novo* TEI rates based on the number of insertions found in their cohort without accounting for detection sensitivity ^6; 20; 21; 49^. Multiple factors, including filtered regions, low sensitivity of the algorithm being used, or false negatives due to the sequencing methodology, result in an underestimate of true *de novo* rates. For example, TEI detection in short read Illumina sequencing data is less sensitive than in long read data, particularly for L1 TEIs^50^. Therefore, we adjusted the observed *de novo* rates to account for sensitivity loss and to obtain precise estimates. Specifically, we measured *xTea* sensitivity on the downsampled (39.4x) Illumina WGS data from HG002, the HapMap sample extensively profiled by multiple sequencing platforms by the Genome in a Bottle consortium^51; 52^ using a high-quality catalogue of haplotype-resolved non-reference TEIs for the sample (see Methods). We obtained sensitivities of 82%, 55%, and 79% for Alu, L1, and SVA respectively and the adjusted *de novo* rates of 1 in 29 births for Alu (95% CI 24-34), 1 in 104 births for L1 (95% CI 77-146), and 1 in 192 births for SVA (95% CI 127-309) (Figure 2A and Table S3). Compared to 1 in 20^3^ or 1 in 21^4^ Alu insertions per birth by earlier studies using evolutionary and mutational based methods, our estimate is lower but within the range from more recent work using family WGS data of 1 in 39.7 births (95% CI 22.4– 79.4)^6^. L1 rates observed here are also within the ranges observed previously of 1 in 63 births (95% CI 30.6–153.8)^6^ and 1 in 95-270^5^ but higher than the Xing *et al.* 2009 rate of 1 in 212 births^4^. Our SVA *de novo* rates are much higher than the Xing *et al.* 2009 rate of 1 in 916 births, but not as high as the Feusier *et al.* 2019 rate of 1:63 births (95% CI 30.6–153.8). Furthermore, the large sample size in our study produced more reliable estimates with smaller confidence intervals than previous analyses (Figure 2A). Recently published work in this ASD cohort^22^ detected fewer insertions and reported 31% (1 in 42 Alu), 55% (1 in 231 L1), and 38% (1 in 309 SVA) lower *de novo* insertion rates than ours, possibly due to their exclusion of mosaic insertions in their rate estimates, the use of a less sensitive pipeline^44^, and not adjusting for the lower sensitivity for detection of TEIs in short read data.

**Figure 2.**
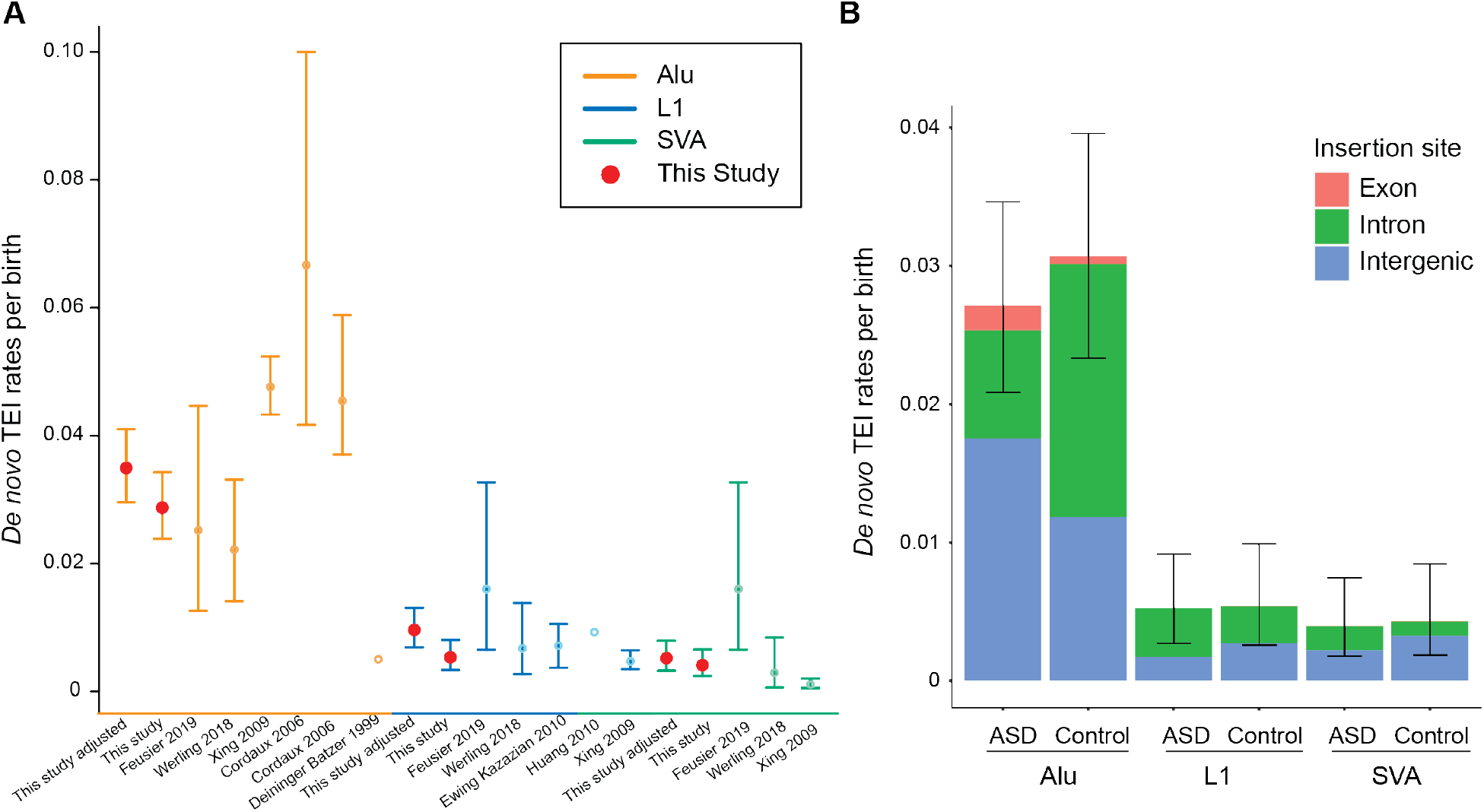
Rates of *de novo* TEIs. **A** Combined rates of *de novo* TEIs per birth for ASD and controls compared to previous studies. The adjusted rate in our study accounts for lower sensitivity for detecting TEIs in short read Illumina data compared to long read sequencing data. **B** Rates of *de novo* TEIs per birth in probands with ASD and unaffected siblings (controls).

We detected 62 *de novo* Alu insertions in ASD (N=2,286) and 57 in controls (N=1,857), 12 *de novo* L1 insertions in ASD (N=2,286) and 10 in controls (N=1,856), and 9 *de novo* SVA insertions in ASD (N=2,288) and 8 in controls (N=1860) (Table S2). We did not detect a difference in total *de novo* TEIs in ASD versus unaffected siblings (Figure 2B) but unexpectedly observed a higher rate of intronic Alu insertions in controls (p=0.003, two-sided Fisher’s Exact Test) (Figure 2B). On the other hand, we observed a trend towards more exonic and intergenic Alu insertions in ASD than controls though not significant (p=0.388 for exonic insertions, p=0.157 for intergenic insertions, two-sided Fisher’s Exact Test) (Figure 2B) which leads to similar overall rates for *de novo* Alu insertions.

We observed *de novo* intronic L1 insertions in syndromic SFARI ASD genes^53^ only in ASD and not in controls, and the rate in ASD was higher than expected (empirical two-sided p-value using 10,000 permutation runs, p=0.001, q-value=0.03) (Figure 3) (Table 1). We also observed a trend for more *de novo* intronic L1 insertions in high pLI genes^54^ in ASD than expected (empirical two-sided p-value, p=0.02, q-value > 0.05) (Figure S4). We observed *de novo* exonic insertions in genes with a high probability of LoF intolerance or haploinsufficiency (pLI ≥ 0.9)^54^ only in affected individuals (Table 1 and Table S2), including an exonic insertion in *CSDE1*, a gene recently implicated in patients with ASD and neurodevelopmental disabilities^55^. There is a large overlap between SFARI genes and high pLI genes with *de novo* L1 insertions in cases; 80% (4/5) of SFARI genes with L1 insertions in ASD are also high pLI genes, suggesting that the *de novo* events can disrupt the haploinsufficient ASD genes and contribute to ASD risk (Table 1).

**Figure 3.**
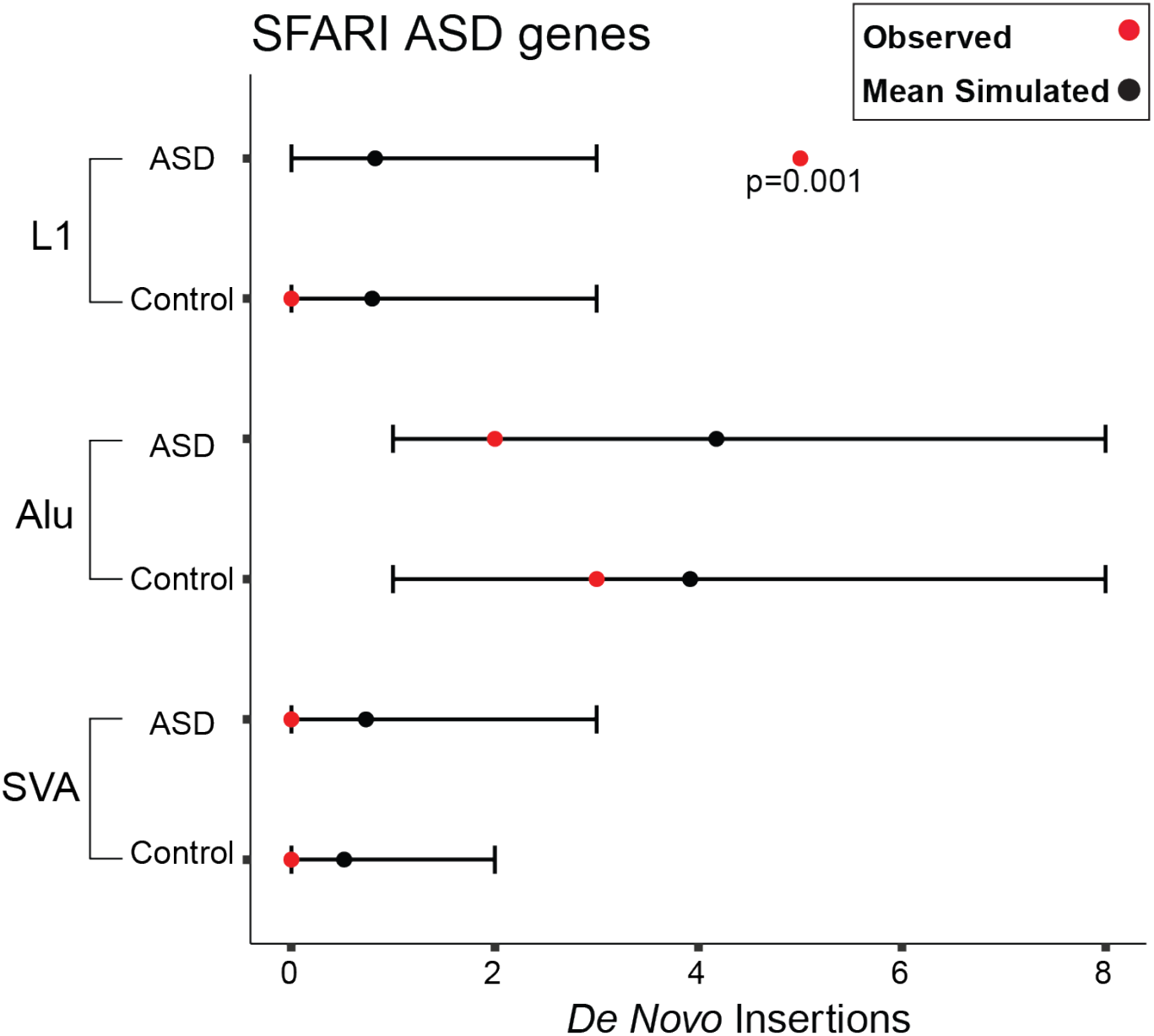
Enrichment of *de novo* TEIs in SFARI ASD genes. Observed numbers of *de novo* TEIs in a list of complied ASD genes are marked by red dots. Black dots and lines represent mean numbers and 95% confidence intervals of expected TEIs based on 10,000 random simulations, respectively. More *de novo* L1 insertions in ASD genes than expected are observed in cases only.

**Table 1:**
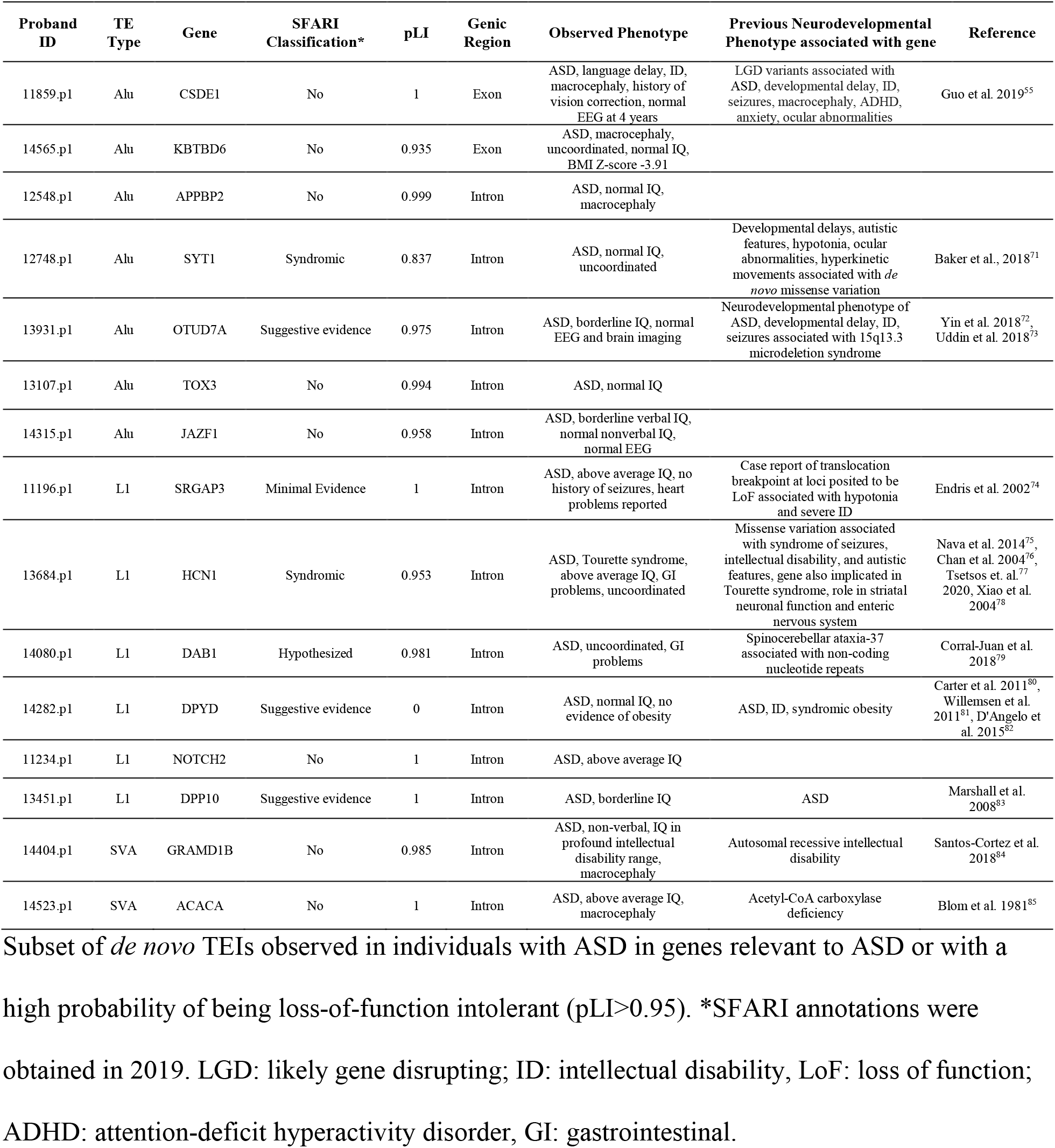
Select *de novo* insertions in ASD and high pLI genes in affected individuals.

Since *de novo* SNVs increase with both paternal and maternal age^56^, and this presents an increased risk to ASD^57–59^, we tested whether there was a difference in parental age at birth in children with and without *de novo* TEIs. We found a modest, but not significant, increase in paternal age for children with *de novo* TEIs compared to those without *de novo* TEIs (M= 33.94, SD= 5.63 *vs.* M=33.29, SD=4.71; t(163.42)=1.4452, p =0.1503) as well as increase in maternal age (M=31.62, SD=4.92 *vs*. M=31.12, SD=4.92; t(163.75)=1.29, p=0.198) (Figure S5). We also estimated the insertion size of polymorphic and *de novo* TEIs by mapping insertion-supporting reads from *xTea* output to TE consensus sequences and obtaining the minimum and maximum mapping coordinates. The distribution of polymorphic L1 insertion size closely resembles previously published data^44^ (Figure S6A). Overall, *de novo* TEIs showed similar size distributions to polymorphic TEIs but showed different patterns from somatic TEIs that showed more severe 5’ truncation^10^ (Figure S6B).

Some genes with *de novo* TEIs in ASD are highly expressed in the brain at all stages of development (Table S4). Using Enrichr^60; 61^, we found an enrichment of *de novo* TEIs in ASD in genes upregulated in the prefrontal cortex, although this was not significant after multiple test correction (p-value=0.0017, Benjamini-Hochberg q-value=0.07), whereas an enrichment was not detected in controls. Additionally, we found that genes with *de novo* TEIs were enriched for calcium-dependent phospholipid binding in ASD (adjusted p-value=0.034) but did not find enrichment for any Gene Ontology terms in controls. Several *de novo* TEIs were also observed in regions with enhancer and promoter chromatin marks in fetal brain development (Table S5). Thus, we evaluated the enrichment of polymorphic and *de novo* TEIs in different genomic and epigenomic regions using the Roadmap Epigenomics 25-state model^62^. Polymorphic L1 and Alu insertions were depleted in exons, enhancers, and promoters (Figure 4; two-sided empirical p<0.0005, Benjamini–Yekutieli q-value<0.0043 for each individual category) whereas SVAs did not show a significant depletion in those regions likely due to the limited number of insertions (Figure S7 and Table S6). *De novo* TEIs overall showed patterns within the expected ranges in most regions, however, we observed a trend for more *de novo* Alu insertions in active enhancer regions in the fetal brain in ASD than expected but not in controls (two-sided empirical p=0.018, Benjamini–Yekutieli q-value=0.3). This suggests the intriguing possibility that Alu insertions in neural enhancers might be a rare cause of ASD, though larger samples sizes are needed to test this.

**Figure 4.**
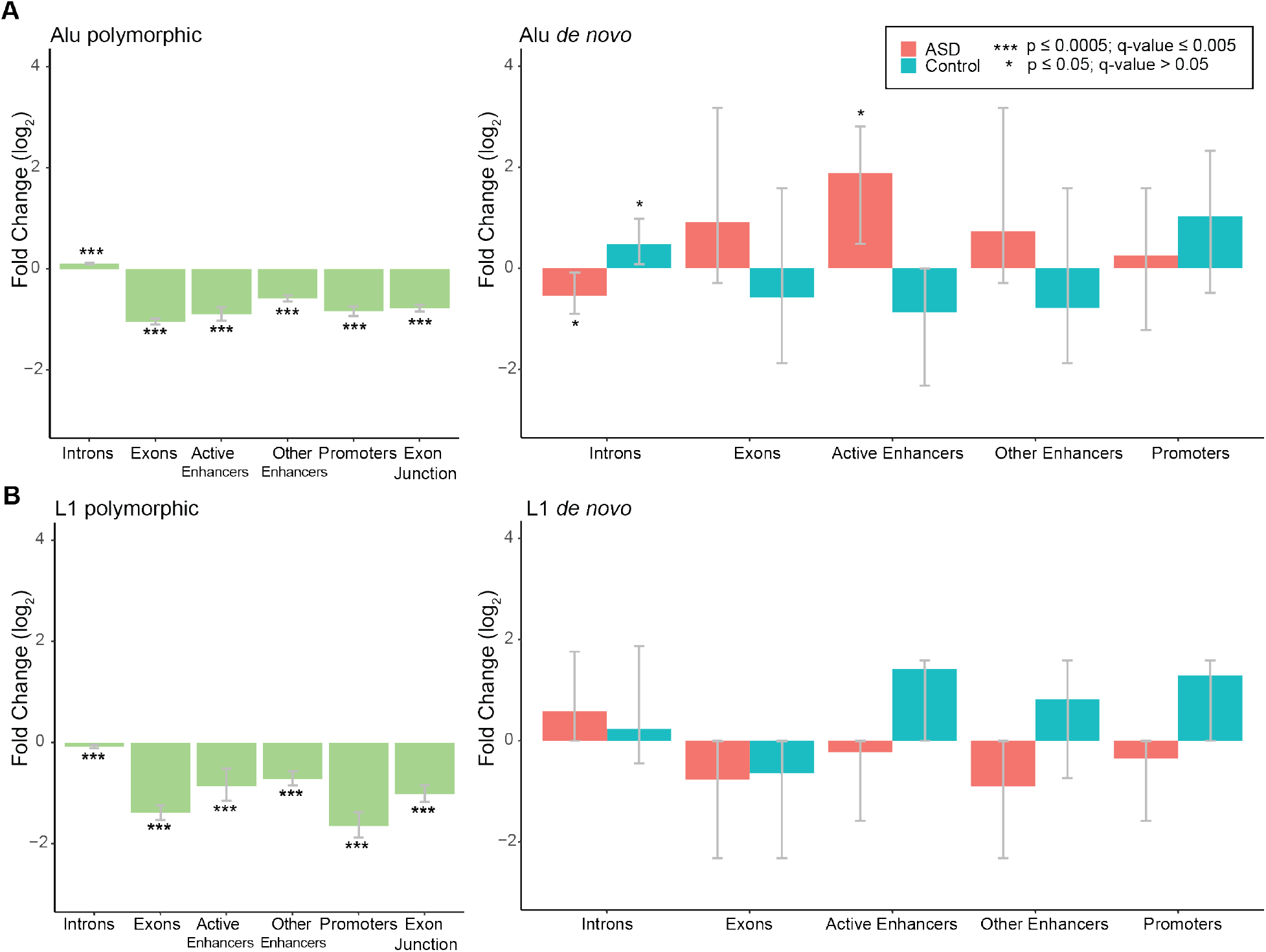
Genomic distribution of polymorphic and *de novo* TEIs. **A** 10,000 random simulations were performed for both polymorphic and *de novo* TEIs based on the observed rates. Log_2_ fold change of observed compared to expected counts in different genomic regions were shown for coding and gene regulatory regions. 95% confidence intervals were estimated based on the empirical distribution of the random simulations. Polymorphic TEIs from parental individuals are depleted in exons and regulatory regions in the developing fetal brain. *De novo* Alu (**A**) and L1 insertions (**B**) do not show this depletion compared to 10,000 random simulations. Two-sided empirical p-values and Benjamini–Yekutieli q-values based on multiple correction of all enrichment and depletions performed are represented.

We selected *de novo* L1 and Alu insertions from both cases and controls in a subset of ASD and high pLI genes as well as in randomly selected genes for full-length PCR validation (Table S7). We validated 22 of 23 (96%) Alu insertions and 6/7 (86%) L1 insertions, achieving a high validation rate of 93% (28/30). Validated insertions include a full-length *de novo* intronic L1 insertion in *DAB1,* a gene with a high probability of being loss-of-function intolerant (pLI=0.981)^54^ and hypothesized ASD gene^53; 63^ implicated in regulating neuronal migration in development via the Reelin pathway in an isoform dependent manner^64^. We additionally validated an exonic Alu insertion in ASD gene *CSDE1^55^* in an ASD proband (Figure 5A). Our manual IGV inspection determined that the exonic Alu insertion in *CSDE1* could either be potentially mosaic at a low allelic fraction in the mother’s blood unless there is low-level contamination from the proband’s DNA, since a single supporting clipped read was observed at the breakpoint (Figure 5B). This insertion was fully validated in lymphoblastoid cell line (LCL) DNA in the individual with ASD and was absent in the mother; LCLs might be limited in validating low-level mosaic mutations (Figure 5A).

**Figure 5.**
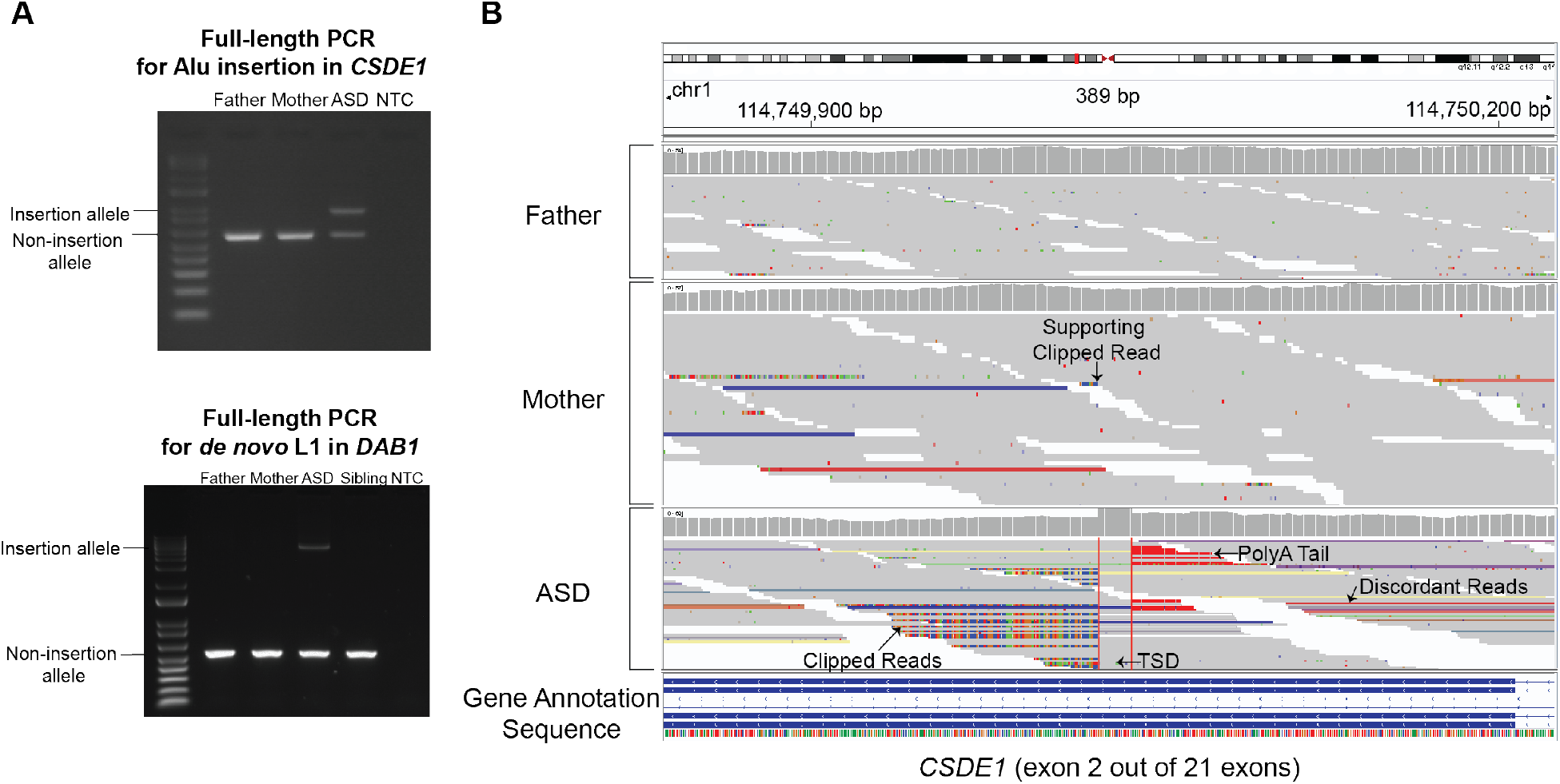
Full-length PCR validations and visual inspection. **A** Full-length PCR validation of the Alu insertion in *CSDE1* and of the *de novo* L1 insertion in *DAB1* in ASD cases. In lymphoblastoid cell line DNA, we validated the insertions in the ASD proband only. NTC: non-template control. **B** Integrative Genomics Viewer image at the insertion site in gene *CSDE1* in an ASD case. For each individual, the sequencing coverage (top) and sequencing reads (bottom) are shown. The insertion shows the canonical signatures of target-primed reverse transcription (TPRT)-mediated retrotransposition: 15bp target site duplication (TSD) between the two insertion breakpoints, a poly-A tail, supporting clipped reads and discordant reads with mates mapping to the consensus Alu sequence. The mother has one small clipped read sequence at the breakpoint which has the same sequence as in the proband, suggesting that the insertion could be mosaic at a low allele frequency in the mother’s blood.

Assigning causality of non-coding mutations based on clinical phenotypes is challenging, given that most known ASD genes have been discovered in the context of coding LoF mutations, yet the majority of individuals with ASD do not have LoF coding mutations identified^29; 31; 65^. In order to understand the clinical phenotypes of individuals with TEIs in high pLI^54^ or known ASD genes^53^, we reviewed the available clinical data and compared this to any known phenotypes associated with the gene, as well as the scientific literature more generally available (Table 1). Exonic insertions are likely to disrupt the coding sequence and are thus of particular interest. We observed one exonic Alu insertion in *CSDE1*, which has been recently associated with ASD^55^. The affected proband shared clinical features, albeit non-specific, consistent with the previously described cohort, including ASD, intellectual disability, macrocephaly, and vision impairment. We additionally observed an exonic Alu insertion in *KBTBD6* (Table 1). Variation in this gene has not yet been associated with a reported neurodevelopmental phenotype that we are aware of. However, *KBTBD6* represents an intriguing candidate gene given its high pLI score (pLI=0.935)^54^ as well as its molecular interactions with known ASD genes *CUL3* and *RAC1^66–69^*. The expression pattern of *KBTBD6* in developing neuronal lineages in the prenatal human cortex also suggests an important role in neurodevelopment^70^. Studying target genes of exonic *de novo* TEIs may shed novel biological insight not captured solely with more commonly studied forms of genetic variation in ASD.

We estimated a rate of underlying exonic TEIs of at least 1 in 2,288 in ASD, which is similar to the rate of 1 in 2,434 cases with developmental disorders reported in a recent whole exome sequencing study^9^. Although this is lower than other types of *de novo* genetic drivers of ASD, such as copy number variation, and the contribution of non-coding variants is thought to be smaller than coding LoF mutations^20^, the strong depletion of polymorphic TEIs in regulatory non-coding regions and enrichment of large *de novo* L1 insertions (~6kb when full-length) in introns of ASD genes in cases but not in control suggest some of these non-coding events may contribute to ASD risk. Since intronic TEIs can affect gene function through various mechanisms, such as altering RNA expression and splicing^1; 8^, TEIs contributing to ASD may present a phenotype different from known phenotypes caused by LoF coding mutations or large CNVs in these genes. Including TEIs and structural variants in standard clinical genetic analyses for ASD will continue to expand our knowledge of non-coding mutations and could increase the rates of genetic diagnoses^17^. Our work also presents important advances in scalable bioinformatic processing and identification of TEIs, which by their nature represent a challenging form of genomic variation to study. Future work, including both further development of computational methods, as well as experimental functional assessment of the effects and pathogenicity of non-coding TEIs, will be critical in understanding the role of these mutations in autism.

## Description of Supplemental Data

Supplemental Data include methods, 7 figures, and 8 tables.

## Supplemental Tables

Table S1. Polymorphic insertions sample sizes

Table S2. *De* novo insertions in ASD and controls

Table S3. *De novo* insertion rates and sample sizes

Table S4. *De novo* insertions which overlap the top 10% expressed genes in the neocortex during development

Table S5. Number of *de novo* insertions overlapping regions with epigenetic annotation in fetal brain

Table S6. Number of observed polymorphic insertions in parental SSC samples overlapping regions with epigenetic annotation in fetal brain

Table S7. PCR validations primers and samples

Table S8. Memory and time cost of xTea on different numbers of CPU cores

## Acknowledgments

We sincerely thank the families who participated in this research in the Simons Foundation Autism Research Initiative (SFARI) and Simons Simplex Collection (SSC). R.B.M was supported by the Fundacion Mexico en Harvard and CONACYT fellowships. C.M.D is supported by the Translational Post-doctoral Training in Neurodevelopment Program (NIMH T32MH112510). E.A.L. is supported by the NIH grants (K01 AG051791, DP2 AG072437), Suh Kyungbae Foundation, and Charles H. Hood foundation. C.A.W is supported by the NIMH (U01MH106883) through the Brain Somatic Mosaicism Network (BSMN). E.A.L and C.A.W are supported by the Allen Frontiers Program through the Allen Discovery Center for Human Brain Evolution. C.A.W is an Investigator of the Howard Hughes Medical Institute. We acknowledge the clinicians who contributed to the collection of samples and phenotypic data, the SSC principal investigators, and the SFARI staff, in particular S. Xiao and R. Rana for providing additional information. This research was performed with credits from the National Institutes of Health (NIH) Cloud Credits Model Pilot, part of the NIH Big Data to Knowledge (BD2K) program. We thank Rutger’s RUCDR Infinite Biologics for providing DNA samples. We are grateful to S. Hill and J. Neil for their help accessing the SSC data, and to M.E. Talkowski, P.R. Loh, E. Macosko, B. Zhao, G.D. Evrony, and R.E Andersen for their advice.

## Data and Code Availability

The most current version of *xTea*, including a filtering and genotyping module used here after running the initial pipeline can be found here: https://github.com/parklab/xTea/. The docker file and version (warbler/xteab:v9) used in our analysis can be found here: https://hub.docker.com/repository/docker/warbler/xteab and the github *xTea* branch used is located here: https://github.com/parklab/xTea/tree/release_xTea_cloud_1.0.0-beta. The code used for detection of *de novo* insertions and for designing primers for full length validation of TEIs is located here: https://github.com/ealeelab/TEs_ASD.

Parental polymorphic insertions classified as filtered high confidence have been submitted to dbVar under the study accession “nstd203”. Intervals for merged insertions within the SSC cohort and cohort parental allele frequencies for these intervals are located here: https://github.com/ealeelab/TEs_ASD/tree/main/SSC_TEs.

