## Supplemental Methods, Figures, and Tables for "Whole-genome analysis of *de novo* and polymorphic retrotransposon insertions in Autism Spectrum Disorder"

### Supplemental Material and Methods

#### Protocol approval and patient consent

This research was performed in accordance to protocols of the institutional review board of the Boston Children's Hospital. Consent to use the data and samples obtained from human samples was obtained from the Simons Foundation Autism Research Initiative.

#### Datasets and data processing with *xTea*

Samples from the Simons Simplex Cohort (SSC) from phases: Pilot, Phase 1, Phase 2, Phase 3-1, Phase 3-1, and Phase 4 were analyzed. These samples are hosted on Amazon Web Services (AWS) and Tibanna[1] was used for managing jobs on AWS. In order to detect germline and *de novo* transposable element insertions (TEIs), we implemented a dockerized version of *xTea* (https://github.com/parklab/xTea) on Amazon Web Services and downloaded the results to perform further filtering with the *xTea* pipeline. For each job, a cloud instance with specific configurations was created, and each individual cram file was downloaded from the S3 bucket to the instance. *xTea* docker was pulled from dockerhub (https://hub.docker.com/repository/docker/warbler/xteab, v9). The reference genome and repeat libraries were also downloaded from the S3 bucket to each instance. *xTea* was run on each downloaded cram, and the results were compressed and saved to the Amazon S3 bucket. The running cost varies based on memory usage and number of cores. We evaluated the performance of *xTea* with different number of cores and memory (Table S8) and used a spot instance with 16 cores and 32G memory (c5.4x large) for all samples.

After removing outlier results and confirming that these were due to corrupted bam files with incomplete sequences or failed *xTea* runs, we analyzed whole genome sequencing data from ~2,288 Autism Spectrum Disorder (ASD) affected individuals and ~1,856 unaffected siblings with both parents sequenced (Table S1 and Table S3 for sample sizes per TE type). The approximate average sample depth, as determined by *xTea*, was 39.4x. Paired-end reads were 151 base pairs in length.

**TEI identification with *xTea***

For each cram file, *xTea* ran three major steps to call TE insertions. First, raw candidate sites were collected based on whether there were enough qualified clipped reads at the breakpoints, where qualified clipped read means part of the read is aligned to the flanking region while the clipped part is well aligned to the consensus TE sequence or other copies. Second, for each passed candidate site we checked whether there were enough discordant reads support. Here, we consider a pair of reads with one read aligned to the flank region and its mate aligned to the TE consensus sequence or other copies as discordant. In addition, intermediate files were saved to record the number of clipped and discordant reads for all the raw candidate sites. Third, we ran TE type specific filters to remove false positives. In specific, while we used the default values for most of the parameters, there are three major parameters (the number of clipped reads, the number of discordant pairs, and the number of clip and discordant reads) which can affect the sensitivity and specificity. We thoroughly evaluated these three parameters and required >= 3 clipped reads, >=5 discordant pairs, and >=1 clip and discordant pairs as the optimal one to maintain a high sensitivity and high specificity. As a consequence of target-primed reverse transcription, a polyA tail and target site duplication should be observed along with enough supporting clipped and discordant reads at both sides of the breakpoint. However, in many cases not all of these features could be detected. *xTea* incorporates a confidence rating system which evaluates whether all these features are found and whether they are on one or both sides of the breakpoint. We selected only insertions classified as “High confidence”. Additional filters within *xTea* include examining the patterns of insertion-supporting clipped sequences and discordant reads mapped to the TE consensus sequences: the supporting reads should not be scattered across the consensus but instead form one cluster (c1) for 5'-clip reads and another cluster (c2) for 3'-clip reads; the mates of 3' and 5' discordant reads should form two distinct clusters (d1 and d2). The distance between c1 and d2 and between c2 and d1 must be less than the average insert size ± 3*(standard-deviation of the insert size). The supplementary *xTea* filter module with TE type specific filters implemented in our analysis is now part of the main *xTea* code in the latest version.

**Polymorphic insertions**

After obtaining the *xTea* filtered high confidence insertions, we excluded calls where the clipped and discordant reads mapped above the consensus size *xTea* uses for mapping for AluY, L1HS, and SVA (282, 6,120, and 1,400 base pairs respectively). This removed some Alu insertions, which tended to be polyA expansion artifacts. Since breakpoint positions can have slight differences between individuals, these insertions were given a 40 base pair margin from the midpoint of the breakpoints and were merged with bedtools[2] "merge" if they overlapped to obtain a unique set of non-redundant TEIs in the SSC cohort. This resulted in 86,154 high confidence filtered unique polymorphic TEIs (68,643 Alu, 12,076 L1, and 5,435 SVA).

**Comparison with known non-reference TEIs**

To determine whether insertions in the SSC cohort were known or novel, merged TEI calls were overlapped using bedtools[2] "intersect" with the breakpoints from gnomAD[3], 1000 genomes[4], or a compilation of other studies[4-12] obtained from Evrony *et al.*[13] to obtain insertions in our cohort that are not found in these studies (novel), known TEIs which overlap, as well as known TEIs which overlap to individual studies only.

To obtain Venn diagrams for overlap with other cohorts, TEIs from unrelated parental individuals were given a 40 base pair margin from the midpoint of the breakpoints and were merged with bedtools[2] "merge" if they overlapped to obtain a unique set of non-redundant TEIs in unrelated individuals in the SSC cohort. Breakpoints from gnomAD[3] and 1000 genomes[4] were also given a 40 base pair margin. The different datasets were overlapped using bedtools[2] “intersect” and counts were plotted in R with the VennDiagram library[14].

##### Allele frequencies and comparison with previous studies

The parental merged insertions described previously were used. These were genotyped with the *xTea* genotyping module which uses a random forest model to genotype TEIs (https://github.com/parklab/xTea). The population allele frequency (PAF) within the cohort was calculated as the number of alleles carrying the TEI in the population divided by the total number of chromosomes in the population. For the comparison with gnomAD PAF, we also estimated PAF within the SSC parental cohort using the Hardy-Weinberg principle. Here, p+q=1, where p is the frequency of the insertion allele in the population and q is the frequency of the non-insertion allele in the population. Assuming that $q^2$ is the fraction of individuals without an insertion, we calculated the PAF as 1 - (sqrt((total parental individuals in the cohort - individuals with insertion allele) / total individuals in the cohort)). Merged breakpoints were overlapped with gnomAD TEIs[3] using a window of 40 base pairs to define overlap. We compared the PAF within the SSC cohort to both the overall PAF and the European PAF in gnomAD since 83% of fathers and 85% of mothers were classified as white (Figure S3).

##### Detection of *de novo* TEIs

We used the high confidence post-filtered insertions from *xTea* for this analysis for Alu, L1, and SVA (https://github.com/parklab/xTea). TEIs were given a 40 base pair margin from the midpoint of the breakpoints and were overlapped with known non-reference (KNR) insertions obtained from previous studies[4-12] as well as reference SVA, reference young L1 (L1HS, L1PA2, L1PA3) or reference young Alu (AluY)[15] and excluded if they overlapped. To exclude inherited insertions which may have been missed in parents, we excluded insertions which had clipped or discordant reads in the raw parental files (clip_reads_tmp0 and discordant_reads_tmp0) in the *xTea* output. We imaged *de novo* candidates on IGV 2.4.19[16] for manual inspection. We visually confirmed the absence of supporting parental reads, as well as the presence of a target site duplication, a polyA tail, and clipped and discordant supporting reads that support a retrotransposition event[17]. Insertions were scored as "high confidence *de novo*" if visual inspection of calls passed these criteria for *de novo* insertions, as "*de novo*" if there were some discordant reads in the parents but no clipped reads supporting the breakpoint in parents, as "somatic candidate" if there is strong read support with a polyA tail, target site duplication, clipped reads and discordant reads, but the supporting reads are a small fraction of the overall coverage at the breakpoint, as "parental mosaic candidate" if there were <=2 clipped reads in one of the parents and discordant reads at a low allele frequency, suggesting it might be mosaic in parent's blood yet not called by *xTea* due to the low allele frequency, and "false negative parental" if it is not clear whether there is a false negative insertion or a mosaic blood insertion in the parental sample due to having few clipped reads but many discordant reads near the insertion site. Only insertions scored as "high confidence *de novo*", "*de novo*", "somatic candidate", and "parental mosaic candidate" were included.

***De novo* retrotransposition rates**

Rates were calculated as the number of *de novo* TEIs for both ASD affected and unaffected siblings divided by the total sample size. Samples which failed the *xTea* run were excluded from the analysis, resulting in a sample size of n=4,142 for L1, n=4,143 for Alu, and n=4,148 for SVA (Table S3). Rates and confidence intervals from previous studies were obtained from Feusier *et al.*[18]. Our 95% confidence intervals were obtained in the same manner, with an exact binomial confidence interval estimate using the binconf R function[19].

With long read technologies, the sensitivity for detection of TEIs is higher[20], suggesting that our raw rates are an underestimate. In order to account for genomic regions in which *xTea* is unable to detect TEIs given the lower sensitivity with Illumina short read data, as well as for the reference filters we used for *de novo* insertions, we calculated our sensitivity for detecting germline TEIs in the Genome in a Bottle sample NA24385/HG002[21] which has been sequenced with both long and short read technologies. A curated set of 9,970 (>50bp) insertions was obtained from Genome in a Bottle V0.6[22] and integrated with 15,268 (>50bp) insertions from a haplotype assembly of the samples[23] (https://github.com/parklab/xTea). RepeatMasker[15] was used to annotate Alu, L1, and SVA sequences, and then insertions were confirmed by manual inspection of poly-A tails and target site duplication or deletions on IGV[24]. This resulted in 1,642 (1,355 Alu, 197 L1 and 90 SVA) high-confidence TEIs detected in the NA24385/HG002 genome but not in the reference genome. We downsampled the HG002 Illumina bam file to the average coverage of SSC samples (39.4x) and detected TEIs with *xTea*. We excluded calls which overlapped reference SVA, reference young L1 (L1HS, L1PA2, L1PA3) or reference young Alu (AluY)[15], as performed for the SSC analysis, and calculated the sensitivity of our pipeline to detect the set of curated TEIs. The sensitivity detected was 82%, 55%, and 79% for Alu, L1, and SVA respectively. We adjusted the number of total ASD and control *de novo* insertions by dividing by the sensitivity and obtained an exact binomial 95% confidence interval using the binconf() R function[19]. Rates between ASD and controls were compared with a 2-sample test for equality of proportions with continuity correction test in R (prop.test)[25].

**Annotation of TEIs**

UCSC Table Browser[26] was used to obtain RefSeq gene annotations and coordinates for exons, and introns. SFARI gene annotations were obtained from SFARI gene[27] in March, 2019. Categories S (Syndromic),1 (High Confidence), 2 (Strong candidate), 3 (Suggestive evidence),

4 (Minimal evidence), and 5 (Hypothesized but untested) were included. *De novo* insertion candidates were also annotated with the probability of being loss-of-function intolerant pLI[28]. Chromatin states from fetal brain tissue (E081 and E082) were downloaded from https://egg2.wustl.edu/roadmap/data/byFileType/chromhmmSegmentations/ChmmModels/imputed12marks/jointModel/final/ [29]. States were classified as: 13_EnhA1 and 12_EnhA2 = active enhancers; 15_EnhAF, 16_EnhW1, 17_EnhW2, 18_EnhAc = other enhancers (weak, flank, acetylation only); 2_PromU, 3_PromD1, 4_PromD2 = promoters. TEIs were overlapped with the two fetal brain regions using bedtools[2] and the number of unique calls in each category was obtained.

**Number of expected insertions in ASD and high pLI genes**

We used a Fisher's exact test to compare the *de novo* rates between individuals with ASD and controls. To test the number of expected insertions in SFARI and high pLI (pLI => 0.9) genes by chance we performed simulation tests. Using the observed *de novo* TEIs in ASD and unaffected siblings, we simulated the same number of random insertions of the same size in the same chromosome with bedtools[2] "shuffle", while also excluding the same young reference regions and KNR regions we excluded when detecting *de novo* calls. We performed 10,000 simulations and determined the number of random insertions which overlapped a SFARI gene, high pLI gene or region of interest per simulation. We determined if the observed value fell on the upper or lower end of the observed distribution to obtain a p-value. A count of 1 was added when estimating this, so for example, the upper p-value=(r+1)/(n+1), where r is the number of simulations greater or equal to the observed value and n is the number of simulations[30]. This value was multiplied by 2 for an empirical two-sided p-value. P-values were corrected for multiple testing with the Benjamini & Yekutieli method[31] using the p.adjust "BY" function in R[25], to account for dependency between tests.

**Mobile Element Insertion Size**

To determine the size of polymorphic and *de novo* insertions, we only included TEIs which had supporting clipped reads on both breakpoints and which did not overlap with reference. Since *xTea* maps reads to several subfamilies of retrotransposons, we excluded calls where the clipped and discordant reads mapped above the consensus size *xTea* uses for mapping for AluY, L1HS, and SVA (282, 6,120, and 1,400 base pairs respectively) since, particularly for Alu calls, these tended to be poly-A expansion artifacts. The consensus sequences within *xTea* were obtained from RepBase23.02[32] and the L1HS consensus sequence was manually constructed by multiple sequence alignment of full length reference sequences. The position of clipped and discordant reads mapping to reference retrotransposon sequences was obtained for each insertion. The minimum position was subtracted from the maximum position to obtain the predicted size. If the maximum position was larger than the consensus length, this was set to the consensus length. The resulting estimated insertion size is an approximation, since we are unable to account for repeat expansions and different poly-A tail lengths[33; 34]. For all insertions including polymorphic and *de novo* TEIs, calls were given a 40 base pair margin from the midpoint of the breakpoints and were merged based on overlap of these coordinates. The median size for all samples with each insertion is reported. Loess regressions were performed in R[25] with a 25% smoothing span.

**Enrichment and depletion of polymorphic and *de novo* TEIs in genomic regions**

Using the observed *de novo* TEIs candidates in ASD and unaffected siblings and the number of unique polymorphic insertions in parents, we simulated the same number of insertions of the same size in random regions of the genome with bedtools[2] "shuffle", while also excluding the same young reference TE regions and KNR regions we excluded when detecting *de novo* calls for *de novo* simulations, and excluding young reference TE regions for polymorphic simulations.

We performed 10,000 simulations to obtain a null distribution and determined the number of calls which overlapped a region of interest per simulation. We determined if the observed value fell on the upper or lower end of the observed distribution to obtain a p-value. A count of 1 was added when estimating this, so for example the upper p-value = (r+1)/(n+1), where r is the number of simulations greater or equal to the observed value and n is the number of simulations[30]. This value was multiplied by 2 for an empirical two-sided p-value. 95% CIs were calculated by obtaining the 0.025 and 0.975 percentiles of the null distribution. The $\log_2$ FC was calculated as the $\log_2$(observed value/mean of the null distribution) and the 95% CIs are plotted as their $\log_2$ values. These p-values along with those for the analysis of estimated insertions in ASD SFARI and high pLI genes were corrected for multiple testing with the Benjamini & Yekutieli method[31] using the p.adjust "BY" function in R[25], to account for dependency between tests.

For overlap of insertions with brain expressed genes, we selected the neocortex regions from Brainspan[35; 36] (ventrolateral prefrontal cortex (VFC), dorsolateral prefrontal cortex (DFC), medial prefrontal cortex (MFC), primary visual cortex (V1C), primary motor cortex (M1C), orbitofrontal cortex (OFC), primary association cortex (A1C), inferior parietal cortex (IPC), primary somatosensory cortex (S1C), superior temporal cortex (STC), inferior temporal cortex (ITC))[37], and obtained the mean expression of each gene per sample in these tissues and then obtained the mean expression per age group for the following categories: Early prenatal: 8-19 PCW, Late prenatal: 21-37 PCW, Childhood (4 months -11 years), Adolescence: 13-19 years, and Adulthood: 21-40 years (Table S4). We then overlapped genes with insertions in ASD and controls with the top 10% of gene expression observed. Using Enrichr[38; 39], we used as the input genes with TEIs in ASD or controls and tested for genes overexpressed in tissues in the Human Gene Atlas list. We also tested for enrichment of gene ontology terms in the subset of genes with TEIs using g:Profiler[40] in only annotated genes, with a user threshold of 0.05 and a significant threshold for multiple testing correction with the g:SCS threshold method.

**Primer design and validation strategy**

24 cases were chosen for validation based on their clinical relevance by selecting mutations that occurred in SFARI, high pLI, or brain expressed genes and 13 cases were randomly selected for a total of 12 L1 insertions and 25 Alu insertions (Table S7). We did not include events overlapping duplicated regions or reference insertions of the same class. A custom pipeline, based on a previously developed pipeline[41], was used to obtain primer sequences for full length validation. Sequences from -800 to -100 and +100 to +800 base pairs from the insertion breakpoint were used to select primers with Primer3[42]. InSilico PCR from UCSC (https://genome.ucsc.edu/cgi-bin/hgPcr) was then implemented to assess whether these primers would amplify a unique region in the genome. Blat[43] (-stepSize=5 -minScore=20 -minIdentity=80) was then used to confirm unique mapping to the genome. If these steps failed, a masked genome (https://hgdownload.cse.ucsc.edu/goldenpath/hg38/bigZips/hg38.fa.masked.gz) was used for the first step.

PCRs were performed using Phusion Hot Start II High-Fidelity DNA Polymerase. Primers were tested and optimized using the Genome in a Bottle sample NA24385/HG002[21], where we had sequencing data and high confidence insertions from the gold standard available. DNA was quantified using a PicoGreen dsDNA quantitation assay before running at least 70 ng of PCR product, when possible, on a 2% agarose gel (for Alu) or a 1% agarose gel and with a Genomic DNA ScreenTape Analysis on an Agilent TapeStation (for L1) for a higher resolution at determining the insertion amplicon size. A 1kb Plus DNA Invitrogen ladder was used. See Table S7 for detailed PCR protocols. Some primer pairs produced additional amplification bands or artifact bands and were further optimized by increasing the annealing temperature and/or decreasing the number of amplification cycles, and some primer pairs produced lower concentrations of DNA and were optimized by increasing the number of amplification cycles and/or decreasing the annealing temperature. If primer pairs did not amplify a unique non-insertion allele and had artifact bands, we did not proceed with validation of those insertions using with those primers. Out of 12 L1 primer pairs designed for validations of *de novo* insertions, we were able to optimize 9 primer pairs, and we optimized 23 Alu primer pairs out of 25. 2 of the L1 primers were selected for mosaic candidates in 1 case and 1 control and were considered separately for validation rates. These 2 cases did not validate in lymphoblastoid cell line DNA.

Validations were performed with 20ng of DNA from each available family member from lymphoblastoid cell lines provided by the Rutgers University Cell and DNA Repository. This was done by confirming the presence of both an insertion and a non-insertion allele band near or at the expected insertion size in the samples with predicted insertions, and only a non-insertion allele band in the other family members. Water was used instead of DNA in the same reaction as a non-template control for each primer pair.

#### Supplemental Figures

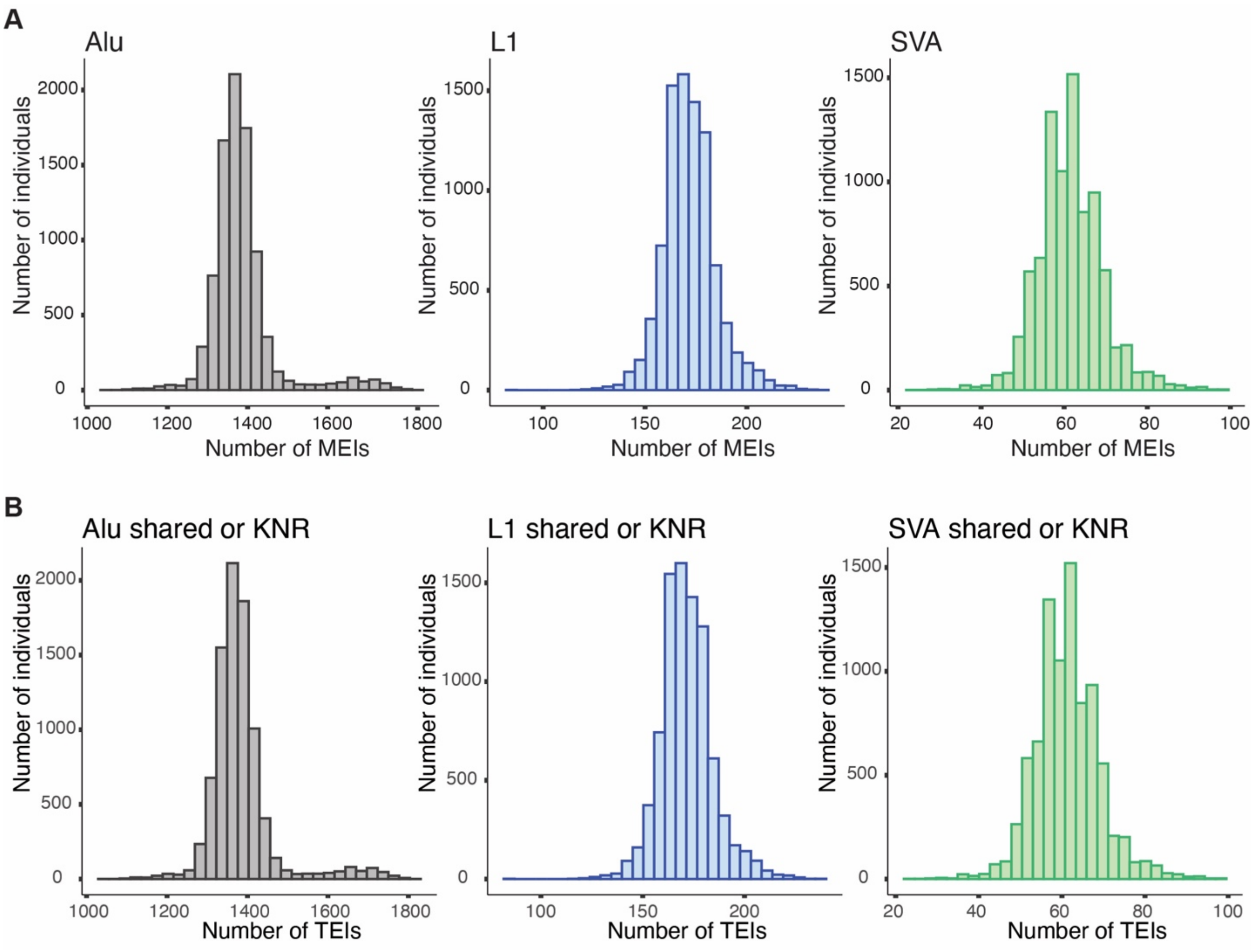


Figure S1. Polymorphic and *de novo* transposable element insertions (TEIs) in the SSC cohort. **A** Number of TEIs detected per individual including parental, ASD, and unaffected siblings. Alu N=8,711, mean=1385.45, SD=82.12; L1 N=8,714, mean=171.94, SD=13; SVA N=8,720, mean=61.50, SD=7.75. **B** Shared polymorphic and known non-reference TEIs in the SSC cohort. Number of TEIs detected per individual including parental, ASD, and unaffected siblings which are found in more than 2 individuals and/or in gnomAD[3] or 1000 genomes[4]. Alu N=8,711, mean=1383.26, SD= 81.23; L1 N=8,714, mean=171.62, SD= 12.89; SVA N=8,720, mean=61.36, SD=7.72.

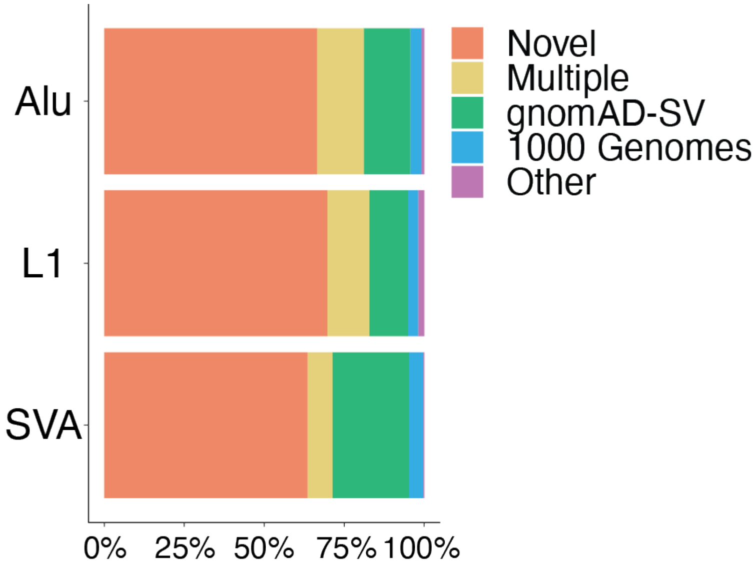


Figure S2. Percentage of insertions which were not found in previous studies (novel) or overlap with TEIs from previous analyses (known) subdivided by source of known overlap[3-12].

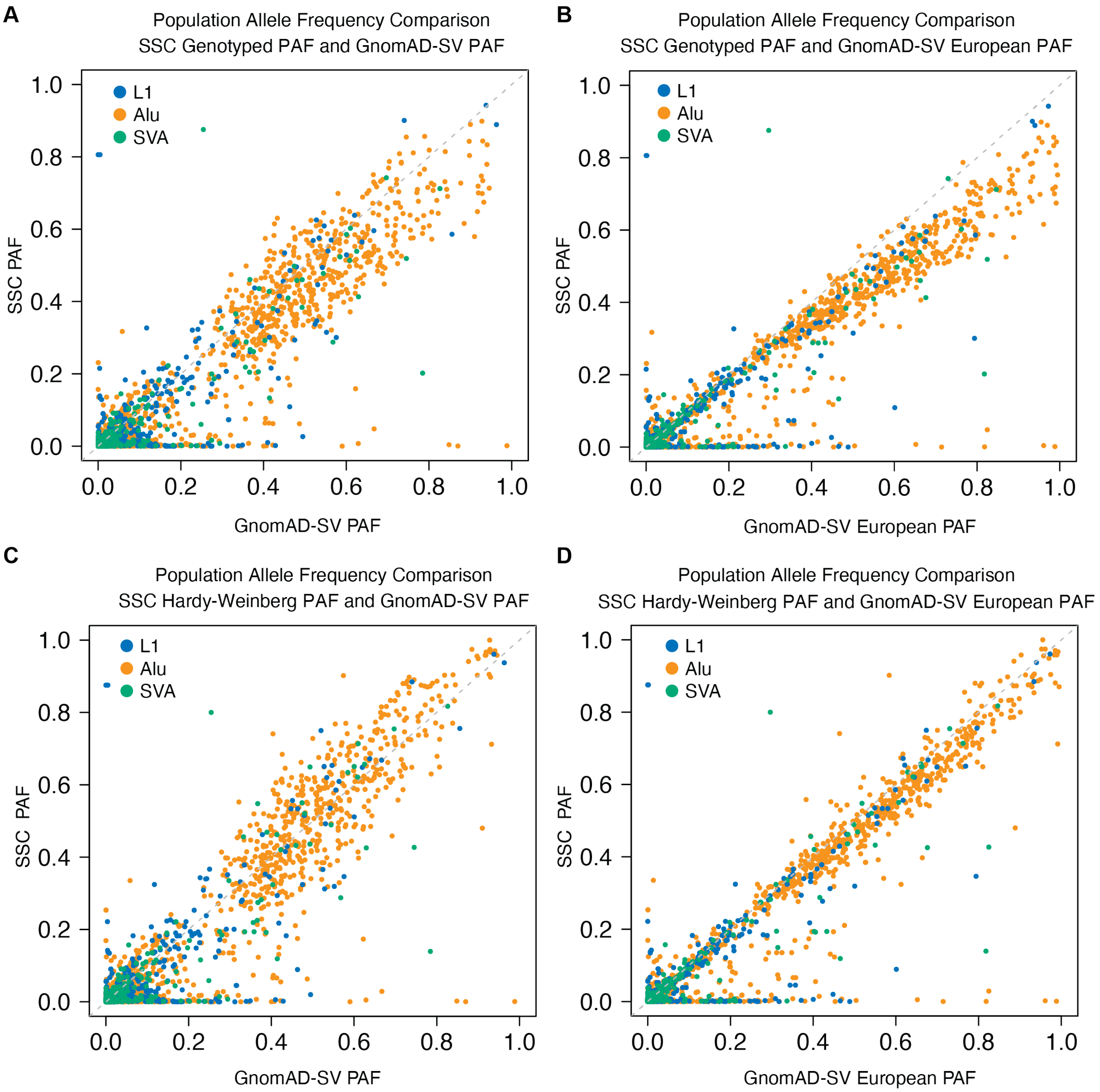


Figure S3. Comparison of population allele frequencies (PAFs) between unrelated parental individuals in the SSC cohort and gnomAD-SV TEIs[3]. **A** Comparison using the SSC PAFs from genotyped TEIs. The PAF was defined as the number of alleles carrying the TEI in the population divided by the total number of chromosomes in the population. GnomAD-SV PAFs are for the entire population or **B** the European population. **C** Same comparison as in Figure S3A with SSC parental PAFs estimated using the Hardy-Weinberg principle compared to gnomAD-SV PAFs in the entire population and **D** gnomAD-SV European PAFs.

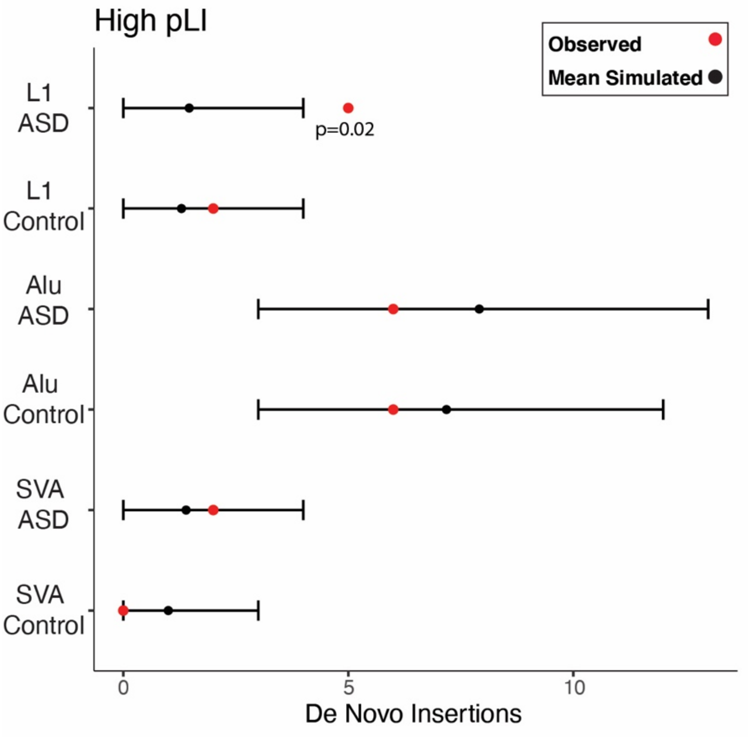


Figure S4. Observed number of *de novo* TEIs in high pLI genes[28] (pLI >=0.90) compared to expected TEIs based on 10,000 random simulations. A trend for L1 insertions in ASD genes than expected are observed in cases (not significant after multiple testing correction with the Benjamini & Yekutieli method).

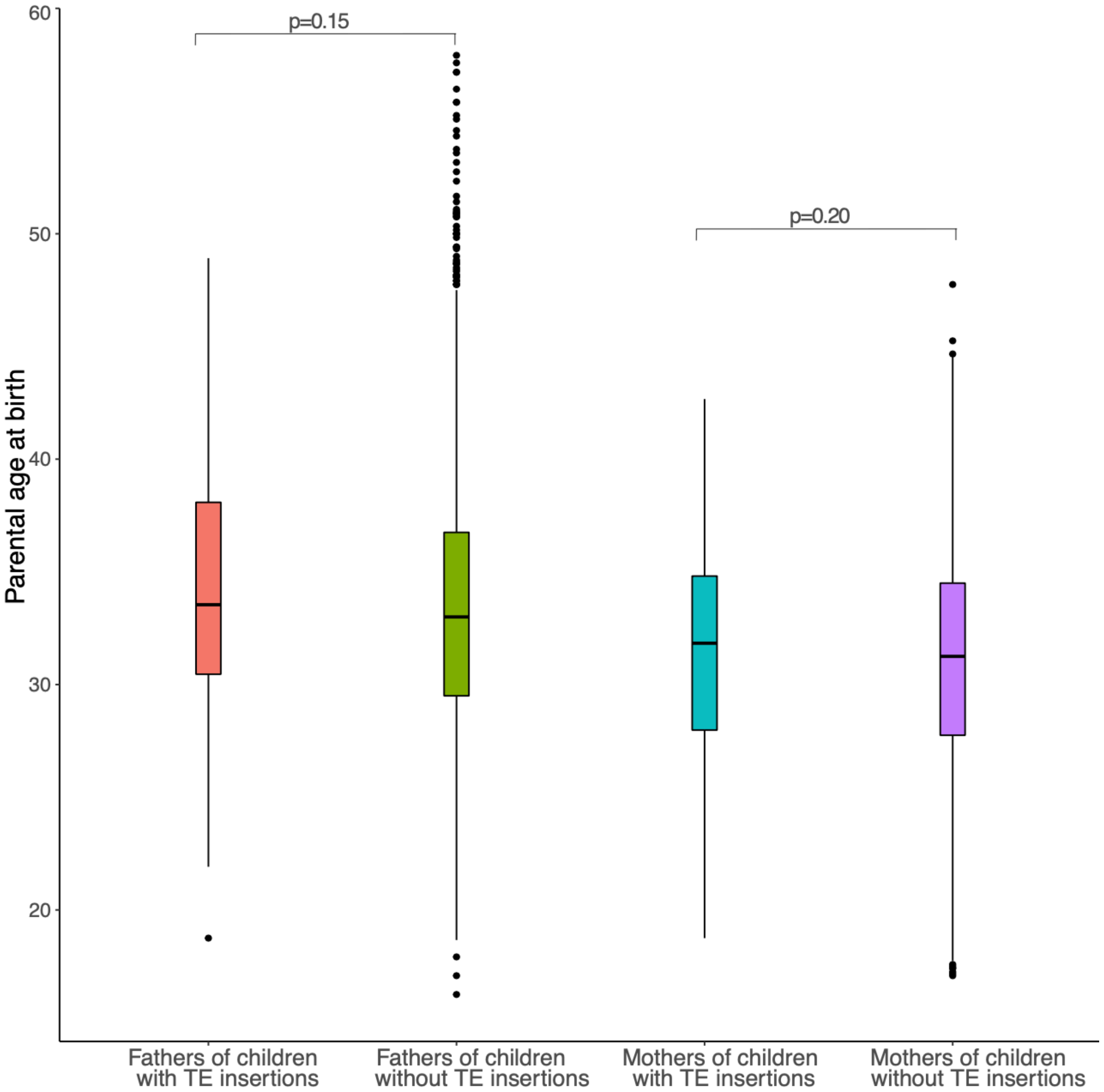


Figure S5. Parental age at birth of children with and without TEIs for cases and controls combined. The median is represented with a line in the middle of the box plot and dots represent outlier samples.

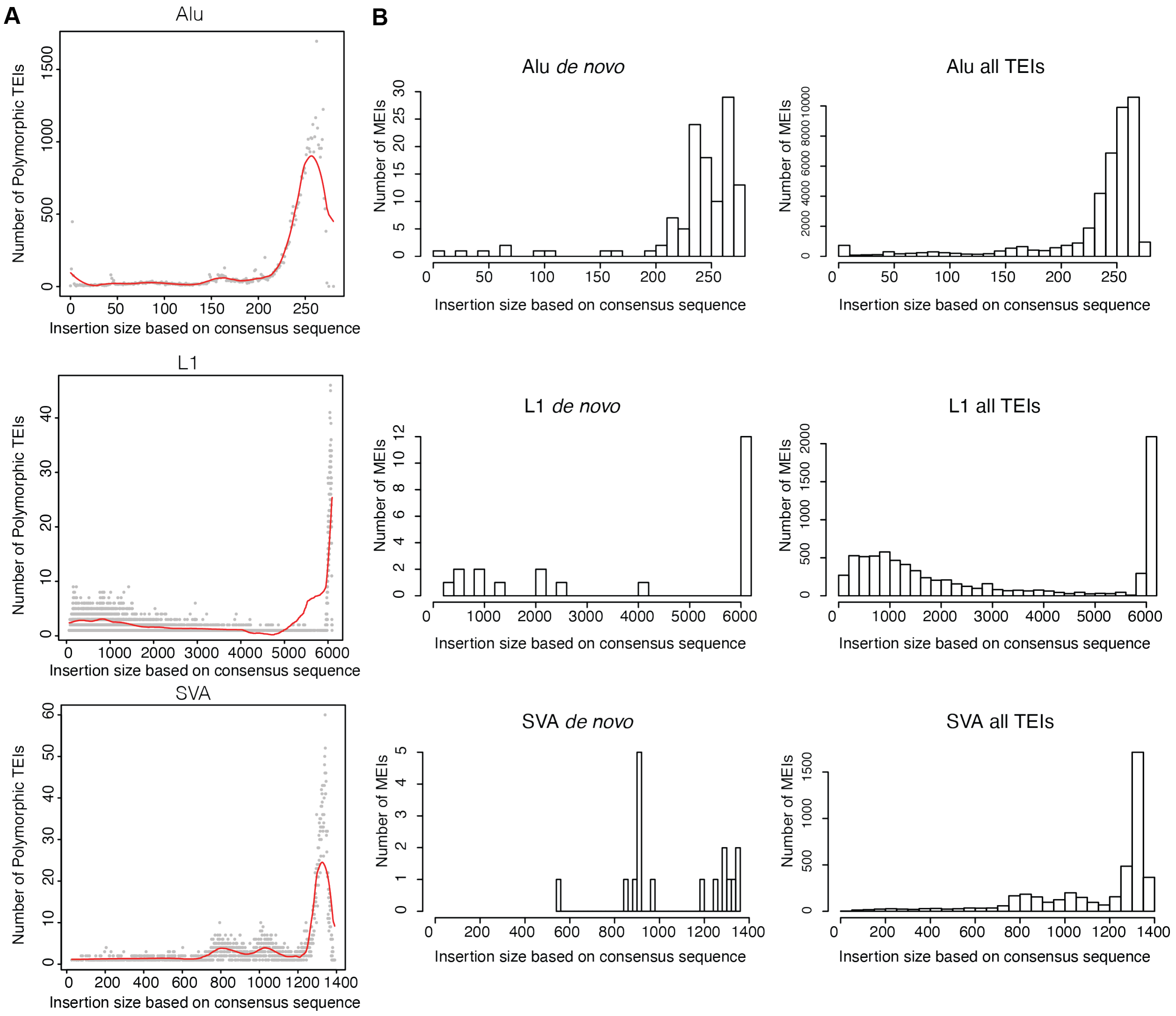


Figure S6. Estimated insertion size of TEIs. **A** Number of TEIs at a certain estimated insertion size for parental, ASD, and control individuals. The red line represents the Loess Regression with 25% smoothing span. Alu N=42,045, L1 N=7,872, SVA N=4,375. **B** Number of TEIs for each insertion size bin for *de novo* insertions or all insertions including both polymorphic and *de novo*. Since we used the position of clipped reads mapping to a consensus sequence to estimate size, this does not account for variable repeat expansions length or polyA tail variability. All: Alu N=42,045, L1 N=7,872, SVA N=4,375; *de novo:* Alu N=188, L1 N=22, SVA N=17.

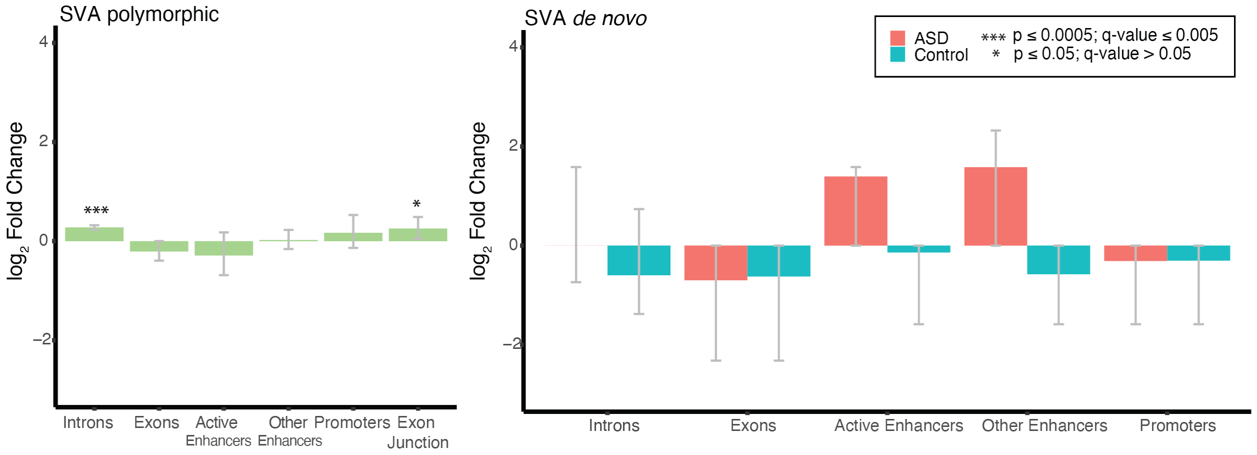


Figure S7. Enrichment and depletion of TEIs in coding and gene regulatory regions. SVA polymorphic and *de novo* TEIs from parental individuals do not show a depletion in exons and regulatory regions in the developing fetal brain, but the number of insertions in these regions are generally fewer than Alu and L1 TEIs (Table S6). 10,000 random simulations were performed for both polymorphic and *de novo* TEIs based on the observed rates. $\text{Log}_2$ fold change of observed compared to expected counts in different genomic regions were shown for coding and gene regulatory regions. 95% confidence intervals were estimated based on the empirical distribution of the random simulations. Two-sided empirical p-values and Benjamini–Yekutieli q-values based on multiple correction of all enrichment and depletions performed are represented.

#### Supplemental Tables

Table S1: Polymorphic insertions sample sizes

| | **L1** | **Alu** | **SVA** |
|---|---|---|---|
| **ASD Cases** | 2,286 | 2,285 | 2,287 |
| **Unaffected Siblings** | 1,856 | 1,855 | 1,858 |
| **Fathers** | 2,286 | 2,285 | 2,287 |
| **Mothers** | 2,286 | 2,286 | 2,288 |

Table S3: *De novo* insertion rates and sample sizes

| | L1 | Alu | SVA |
|---|---|---|---|
| **ASD sample size** | 2,286 | 2,286 | 2,288 |
| **Controls sample size** | 1,856 | 1,857 | 1,860 |
| **ASD number of *de novo* insertions** | 12 | 62 | 9 |
| **Controls number of de novo insertions** | 10 | 57 | 8 |
| **ASD *de novo* rates** | 0.0052 | 0.0271 | 0.0039 |
| **Control de novo rates** | 0.0054 | 0.0307 | 0.0043 |
| **ASD + Control *de novo* insertions** | 22 | 119 | 17 |
| **ASD + Control sample size** | 4142 | 4143 | 4148 |
| **ASD + Control *de novo* rates** | 0.0053 | 0.0287 | 0.0041 |
| **1 in X births** | 188.27 | 34.82 | 244.00 |
| **Sensitivity 39.4x HG002** | 0.553299 | 0.82214 | 0.788889 |
| ***De novo* rates adjusted** | 0.0096 | 0.0349 | 0.0052 |
| **1 in X births adjusted** | 104.17 | 28.62 | 192.49 |
| **Confidence interval rates lower adjusted** | 0.0069 | 0.0296 | 0.0032 |
| **Confidence interval rates upper adjusted** | 0.0131 | 0.0410 | 0.0079 |
| **Confidence interval lower 1 in x adjusted** | 145.79 | 33.84 | 308.56 |
| **Confidence interval upper 1 in x adjusted** | 76.56 | 24.40 | 126.77 |

Table S4: *De novo* insertions which overlap the top 10% expressed genes in the neocortex during development

| | Early Prenatal | Late Prenatal | Childhood | Adolescence | Adulthood |
|---|---|---|---|---|---|
| **Genes with *de novo* Alu insertions in ASD** | CSDE1<br>SYT1<br>KBTBD6<br>TCF25<br>EPS15 | CSDE1<br>SYT1<br>TCF25<br>RPH3A<br>EPS15 | CSDE1<br>SYT1<br>TCF25<br>RPH3A<br>EPS15 | CSDE1<br>SYT1<br>TCF25<br>RPH3A<br>EPS15 | CSDE1<br>SYT1<br>TCF25<br>RPH3A<br>EPS15 |
| **Genes with *de novo* Alu insertions in control** | DCLK2<br>SF3A1 | DCLK2 | DCLK2 | DCLK2 | |
| **Genes with *de novo* L1 insertions in ASD** | DAB1 | | | | |
| **Genes with *de novo* L1 insertions in control** | EPHA7 | | | | |

Overlap of genes with *de novo* insertions and the top 10% expressed genes in neocortex brain regions during development. Early prenatal: 8-19 postconceptional weeks (PCW), Late prenatal: 21-37 PCW, Childhood (4 months -11 years), Adolescence: 13-19 years, and Adulthood: 21-40 years. Genes with *de novo* SVA insertions did not overlap with any of the categories.

Table S5: Number of *de novo* insertions overlapping regions with epigenetic annotation in fetal brain

| Sample and Genomic Region | Alu | L1 | SVA |
|---|---|---|---|
| **ASD introns** | 18 | 8 | 4 |
| **Control introns** | 34 | 5 | 2 |
| **ASD exons** | 4 | 0 | 0 |
| **Control Exons** | 1 | 0 | 0 |
| **ASD active enhancers** | 3 | 0 | 1 |
| **Control active enhancers** | 0 | 1 | 0 |
| **ASD other enhancers** | 4 | 0 | 2 |
| **Control other enhancers** | 1 | 1 | 0 |
| **ASD promoters** | 1 | 0 | 0 |
| **Control promoters** | 2 | 1 | 0 |

Table S6: Number of observed polymorphic insertions in parental SSC samples overlapping regions with epigenetic annotation in fetal brain

| Genomic Region | Alu | L1 | SVA |
|---|---|---|---|
| **Introns** | 28,498 | 4,564 | 2,727 |
| **Exons** | 1,065 | 140 | 169 |
| **Active enhancers** | 254 | 44 | 34 |
| **Other enhancers** | 1,560 | 249 | 205 |
| **Promoters** | 449 | 42 | 81 |
| **Exon junctions** | 1,031 | 145 | 188 |

Table S8. Memory and time cost of xTea on different numbers of CPU cores

| Cores | Memory | Time |
|---|---|---|
| 1 core | 3,011,224K | 30hrs |
| 2 cores | 5,395,264K | 22hrs |
| 4 cores | 7,575,888K | 16hrs |
| 8 cores | 15,112,408K | 7hrs |
| 16 cores | 19,256,964K | 3hr 50mins |
